# Validation of a refined protocol for mouse oral glucose tolerance testing without gavage

**DOI:** 10.1101/2024.09.13.612859

**Authors:** Katherine R. Pye, Louise Lantier, Julio E. Ayala, Craig Beall, Kate L.J. Ellacott

## Abstract

A glucose tolerance test (GTT) is routinely used to assess glucose homeostasis in clinical settings and in preclinical research studies using rodent models. The procedure assesses the ability of the body to clear glucose from the blood in a defined time after a bolus dose. In the human clinical setting, glucose is ingested via voluntary consumption of a glucose-sweetened drink. Typically, in the rodent GTT oral gavage (gavage-oGTT) or (more commonly) intraperitoneal injection (IPGTT) are used to administer the glucose bolus. Although used less frequently, likely due to investigator technical and experience barriers, the former is the more physiologically relevant as it integrates the gastrointestinal tract (GI), including release of key incretin hormones. However, orally gavaging glucose in the GTT is also not without its limitations: gavaging glucose straight into the stomach bypasses potentially critical early glucose-sensing via the mouth (cephalic phase) and associated physiological responses. Furthermore, gavaging is stressful on mice, and this by itself can increase blood glucose levels. We have developed and validated a refined protocol for mouse oral GTT which uses a voluntary oral glucose dosing method, micropipette-guided drug administration (MDA), without the need for water deprivation. This approach is simple and non-invasive. It is less stressful for the mice, as evidenced by lower circulating corticosterone levels 10 minutes after glucose-dosing compared to oral gavage. This is significant for animal and investigator welfare, and importantly minimising the confounding effect of stress on mouse glucose homeostasis. Using a randomised cross-over design, we have validated the MDA approach in the oGTT against oral gavage in male and female C57BL/6J and C57BL/6N mice. We show the ability of this method to detect changes in glucose tolerance in diet-induced obese animals. Compared to oral gavage there was lower inter-animal variation in the MDA-oGTT. In addition to being more representative of the human procedure, the MDA-oGTT is easy and has lower barriers to adoption than the gavage oGTT as it is non-invasive and requires no specialist equipment or operator training. The MDA-oGTT a more clinically representative, accessible, and refined replacement for the gavage-oGTT for mouse metabolic phenotyping, which is simple yet overcomes significant deficiencies in the current standard experimental approaches.

## Introduction

Metabolic diseases, such as type-2 diabetes, are widespread debilitating conditions that often affect the sufferer throughout their life^1^. A common feature of many of these diseases are defects in maintaining glucose homeostasis. The clinical test for glucose intolerance – as a proxy measure for insulin resistance – in people is the oral glucose tolerance test (oGTT), where the subject is given a glucose bolus dose in the form of a sugary drink, and their blood glucose is measured before and at usually two hours after, to assess how well they clear this extra glucose from their system.

In pre-clinical research, the GTT is routinely carried out in rodent metabolic phenotyping studies to assess new treatments for metabolic disorders and gain novel understanding of underlying physiology and pathophysiology^2^. However, the procedure in rodents differs significantly from that carried out in the clinical setting. A key difference is the method of glucose dosing, which is typically carried out either via intraperitoneal (IP) injection or oral gavage. The former bypasses the entire gastrointestinal (GI) tract, and therefore does not incorporate the fundamental physiological responses mediated by this system such as release of key incretin hormones^3^. Although the oral gavage method does include the GI factors, it bypasses crucial cephalic responses^4^.

Despite the oral gavage being more physiologically representative than the use of an intraperitoneal injection, the latter is more widely used in rodent GTTs based on a sampling of papers published in 2007 (73 of 100 studies)^5^ or 2021 (45 of 93 studies)^3^. This is likely due to knowledge and skills gaps relating to oral gavage, which is potentially difficult to master without significant training and experience. A further implication of using these methods is that dosing a rodent by either IP injection or gavage is invasive and requires some form of restraint, which, particularly in mice, generates a significant stress response. Indeed, in the recent comparative study of Small and colleagues, despite the differences in glucose absorption and peak blood glucose between the IPGTT and gavage-oGTT, there was an equivalent elevation in the plasma corticosterone levels (compared to baseline in naïve animals)^3^. This implies that changes in this stress hormone are likely related to the procedural elements, such as handling and dosing, as opposed to being solely glucose-induced. However, stress can affect glucose metabolism^6^, so the stress associated with these tests might introduce variance to the data that could otherwise be mitigated and in some cases may mask physiological effects. Stress-associated data variance may result in the need for a larger sample size for adequate power, leading to increased numbers of animals being used.

Alternatives to gavage for oral dosing for drug-delivery have already been explored, including micropipette-guided drug administration (MDA). Scarborough *et al.* and Schalbetter *et al.* developed this as a method of oral dosing mice with a condensed milk solution containing risperidone or clozapine-*N*-oxide (CNO; for the activation of designer receptors exclusively activated by designer drugs [DREADDs]), to avoid both the stressor of an IP injection and the poorly controlled dosing of drinking water^7,8^. This method has also recently been adapted by an independent research team for tamoxifen administration^9^.

Adapted from the published MDA protocol, herein we present a validated, simple, effective and non-invasive method of voluntary oral glucose dosing for use in the oGTT.

## Methods

### Mice

All animal studies were conducted in accordance with the UK Animals in Scientific Procedures Act 1986 (ASPA) and were approved by the institutional Animal Welfare and Ethical Review Body at the University of Exeter.

C57BL/6J and C57BL/6N mice of both sexes were either ordered from Charles River (Margate, UK) or bred in-house at the University of Exeter (C57BL/6J only) from mice purchased from Charles River. Mice were 8 weeks old at the start of experiments. Unless stated otherwise, mice were housed in same-sex groups of four in standard individually ventilated cages (IVCs), with *ad libitum* access to food (standard laboratory rodent diet [LabDiet (EU) Rodent diet 5LF2; LabDiet, London, UK]), water, enrichment (wooden chew block or ball) and hideaway, on a 12:12 hour light cycle (lights on 0700-1900), at 22 ± 2°C. Cages were changed once a week, but never on the day of or the day prior to an experimental procedure. When cages were changed, some bedding and enrichment from the previous cage was moved with the mice to reduce stress. For high-fat diet (HFD) studies C57BL/6J mice had *ad libitum* access to high-fat diet (Rodent diet with 60% fat [D12492i]; Research Diets, Inc, New Brunswick, USA) for 16 weeks as their only source of food. During the HFD period, mice were separated if they showed signs of overgrooming cage-mates and were weighed and monitored twice a week for adverse reactions to the HFD (e.g., oily fur or irritation from overgrooming).

For experiments carried out at Vanderbilt MMPC-Live, experimental procedures were approved by the Institutional Animal Care and Use Committee at Vanderbilt University. 8-week-old mice of both sexes were ordered from The Jackson Laboratory (Bar Harbor, ME). Mice were housed in same-sex groups of four in Allentown IVCs with *ad libitum* access to food (standard laboratory diet [Picolab® Rodent diet 5L0D; LabDiet (US)]) and water, on a 12:12 hour light cycle, at 21 ± 1.1°C. Cages were changed once a week.

### Handling habituation

Mice were handled using tube or cupping methods throughout the studies^10–12^. Mice were handled daily for 3-4 days prior to the start of MDA habituation by the same handler (female). Habituation included gentle handling above the home cage for approximately 1 minute per mouse each day. The same handler carried out all experiments at the University of Exeter, and handling habituation occurred in the same room as the GTT or other dosing procedure (such as for terminal blood collection). For experiments at Vanderbilt, a different handler may have performed the habituation and the experiment (all female). Mice that were due to be gavaged were also scruffed each day prior to the experimental day for habituation to the gavage procedure, whilst the MDA-group mice underwent MDA habituation.

### Micropipette-guided dosing (MDA) habituation

Mice were habituated to MDA-glucose over a period of 6 days. A 40% glucose solution (in water) was used for the habituation period, with chocolate flavouring (N!ck’s stevia drops – chocolate flavour) at the ratio of 3 drops per 5ml glucose solution. The process was developed from the published habituation method^7,8^, but broadly is as follows: day 1, mice were restrained via scruff and the flavoured glucose solution was presented with the pipette at a horizonal angle; day 2, mice were gently restrained on the food hopper with a light grip on their tail and above their hind legs, and the flavoured glucose solution was presented as before; day 3, mice were allowed to drink from the pipette without restraint, or with a gentle hold on their tail if they were particularly mobile. If they did not drink freely or with the gentle tail restraint, they were restrained again as on day 2. Habituation continued in this manner, with mice first being offered the flavoured glucose solution without any restraint, and restraint only used after a minute of no interest in pipette. See **Supplementary Figure 1** for step-by-step pictures.

### Oral Glucose Tolerance Tests (oGTT)

A within-subjects randomised cross-over design, with route of administration (RoA) balanced between groups, was used to compare oGTT profiles from animals administered oral glucose via MDA and gavage routes. For each test, mice had food removed at lights-on (7:00am) 4 hours prior to the oGTT at 11:00am but had *ad libitum* access to water. Mice were given a bolus dose of 2.5g glucose/kg body weight, flavoured with chocolate stevia drops (N!ck’s) at 3 drops per 5ml glucose solution, via either MDA (80% glucose solution in water) or oral gavage (40% glucose solution in water), where a dose for a 20g mouse measured 62.5µl or 125µl for MDA and gavage, respectively. Animals were randomly assigned to which route of glucose administration they would receive first prior to the habituation period via the out-of-a-hat method, with equal numbers of each sex being assigned to each group and the order of route of administration balanced between groups. Blinding of the investigator for the GTT was not possible as the route of administration was the variable. Immediately prior to the glucose bolus dose (time 0), and at 15, 30, 45, 60, 90 and 120 minutes after, blood glucose was measured via a handheld glucometer (Accu-chek Performer, Roche, with Inform II strips) from a small drop of blood (1µl) gained from the tip of the tail via needle-prick. EMLA cream (Williams Medical Supplies) was used as a local anaesthetic, applied to the tails 30 minutes prior to basal blood glucose measurement. Animals were given at least 1-week between oGTTs. Dosing and blood glucose measurement occurred atop the hopper of a clean cage, separate and out of sight of the home cage.

For the GTTs carried out at Vanderbilt MMPC-Live, mice were handled and housed as described previously, although three handlers were involved throughout the study due to time constraints. Blood was sampled via a minor excision at the tip of the tail, and larger blood samples (40µl) were taken 15 minutes prior to dosing, and 15 minutes after for measurement of plasma insulin. It should also be noted that within this study, one handler was responsible for dosing all females, whilst the other two dosed all males. Blood glucose at Vanderbilt MMPC-Live was measured via a Bayer Contour Next glucometer.

### Plasma corticosterone assessment

C57BL/6J mice of both sexes (11 male, 7 female) were assigned randomly to either the MDA or gavage group via the out-of-a-hat method, with as close to equal numbers of each sex being assigned to each group as possible. MDA-group mice were habituated to MDA-glucose as described, whilst the gavage group was habituated to the gavage procedure via daily handling and gentle scruffing. On the day of procedure, mice were given 2.5g glucose/kg body weight of flavoured glucose solution, then ten minutes later they were euthanised via cervical dislocation and trunk blood collected into a tube containing a drop of EDTA (Invitrogen UltraPure 0.5M EDTA) for a final concentration of 1.5mg EDTA/ml blood). All procedures were carried out in the same 90-minute lights-on period between 10:00-11:30am, over consecutive days. Plasma was isolated from the terminal blood collections via centrifugation (3000rpm for 15 minutes at 4°C) and was run in duplicate on a competitive corticosterone ELISA (Cayman Chemicals #501320) after a single freeze-thaw, at a 1/100 dilution in ELISA buffer. Data were analysed via the Cayman Chemical ELISADouble analysis tool. During the ELISA, the investigator was blinded to the groups that the plasma samples were derived from.

### Plasma insulin assessment

Blood samples taken before and after the glucose bolus within the GTT at Vanderbilt MMPC-Live were spun via centrifugation at 13,000*g* for 1 minute at 4°C to separate the plasma. Plasma samples were run in duplicate on a rat insulin radioimmunoassay (RIA) (Merck #RI-13K) at the Vanderbilt Analytical Services Core.

### Statistical analysis

Data were processed using Microsoft Excel 2013. All values were collated in Prism 10 (GraphPad, San Diego, CA) for tests of statistical significance. The threshold for statistical significance was set at p<0.05. The experimental unit was the individual mouse.

For between-subjects design with one value per subject (e.g., corticosterone data), unpaired t-tests were used. For within-subjects design with two values per subject, paired t-tests were used (e.g., AUC data for MDA-oGTT vs gavage-oGTT). AUC data were corrected for baseline to account for any statistically different basal blood glucose measurements (i.e., time 0 in the GTT). For between-subject statistical comparisons with multiple values per subject, two-way ANOVA was used with Fisher’s LSD post-hoc test applied to compare between column means. For within-subjects design with two values per subject over time (e.g., basal-blood glucose before and after HFD within sub-strain), two-way repeated measures ANOVA was used with Fisher’s LSD post-hoc test applied to compare between column means. Mixed-effects models were used when there were any missing data points. For within-subjects design with three or more values per subject over time (e.g., glucose clearance curves over time for MDA-oGTT vs gavage-oGTT within sub-strain), two-way repeated measures ANOVA was used with Greenhouse-Geisser correction and Šidák’s post-hoc test applied to compare between column means. For within-subjects design with three or more values per subject over time measuring the effect of three independent variables (e.g., glucose clearance curves over time for MDA-oGTT vs gavage-oGTT, male vs female within sub-strain), three-way repeated measures ANOVA was used with Greenhouse-Geisser correction and Tukey’s post-hoc test applied to compare between column means. Male and female mice were used for experiments and data were pooled when two- or three-way ANOVA (as applicable) did not reveal a statistically significant effect of sex or a statistically significant interaction of sex and another variable **[Supplementary Tables 1-3b]**. Graphs were generated in Prism 10. All data are presented as mean ± SEM unless otherwise stated.

Two mice were excluded from the study due to gavage injury, and one mouse was excluded after displaying excess grooming leading to injury within the HFD study.

## Results

### Mice habituated readily to micropipette-guided dosing (MDA) of flavoured glucose

To our knowledge, all published MDA studies to date^7,8^ have used sweetened condensed milk as a delivery vehicle. Due to having a mixed macronutrient composition, sweetened condensed milk was not ideal for the oGTT so we replaced this with a glucose solution. In developing the method, we had initially expected that the mice would be willing to drink a straight glucose solution due to its sweetness; however, mice did not seem to find the offered solution enticing and most failed to habituate to drinking it as expected, despite us successfully habituating animals to drinking sweetened condensed milk (*data not shown*). This is likely due to the low vapour pressure of glucose^4^, meaning that it does not produce a strong olfactory stimulus which may impact the time that it takes to form an effective association.

Next, we tried flavouring the glucose solution with chocolate stevia drops, as used in Kennard et al., 2022^13^, to integrate effective olfactory and flavour stimulus. This worked well for MDA-habituation and was used for all future experiments.

Mice were habituated to the flavoured glucose over 6 days. Across all Exeter studies, by day 6, of 67 mice (C57BL/6J and C57BL/6N mice of both sexes) habituated to MDA, only 2/67 (2.98%) required gentle holding atop the hopper before drinking the solution, while 37/67 (55.22%) drank immediately without any restraint, and 28/67 (41.79%) only required a very gentle tail hold for them to remain in place to drink the solution [**Figure 1**]. Where mice experienced a randomised crossover, only the habituation data from those that had MDA habituation in the first week were included in this analysis (24 mice of the 67 in this figure) to remove any influence of the additional handling. On the day of the procedure, all mice drank the full glucose dose within 30 seconds of presentation.

**Figure 1:**
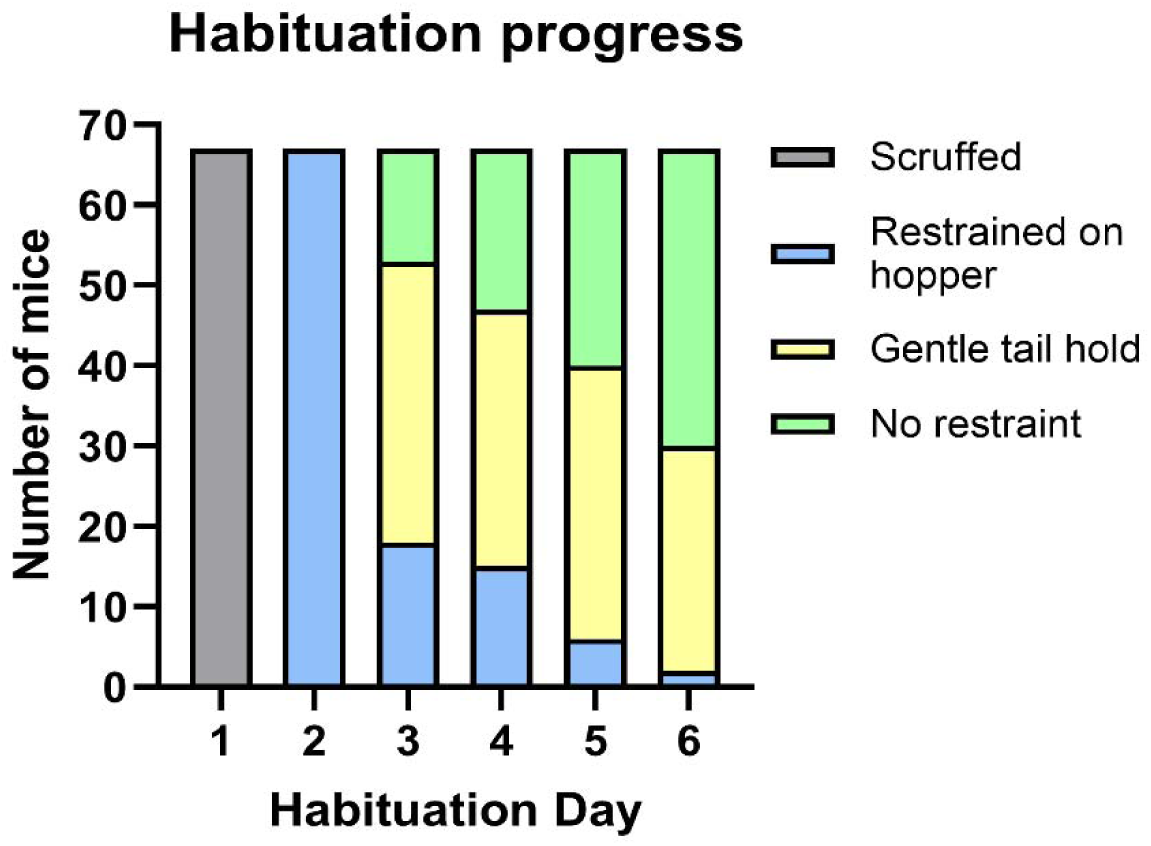
Summary of the progression of mice through the stages of micropipette-guided drug administration (MDA) habituation. Graph shows progress during first-time habituation of all male and female C57BL/6J and C57BL/6N mice from the studies conducted in Exeter. N=67 mice.

### Plasma corticosterone levels were lower in mice dosed with flavoured-glucose via MDA-dosing than via gavage

Activation of the stress response can introduce significant variation in physiological assays. Critically for assessments of glucose homeostasis, stress can increase circulating glucose, including via the action of the hormone corticosterone^14^. Therefore, reducing the impact of stress, a confounding variable within a procedure, is beneficial to not only the experience of the animal, but also the quality of the data.

To assess relative procedural stress, mice were given the same flavoured glucose solution via either MDA or gavage, and corticosterone was measured from their terminal blood plasma 10 minutes after dosing. In gavage-dosed mice plasma corticosterone was significantly higher compared to MDA-dosed mice (94.64 ± 18.63 ng/ml [MDA] vs 233.6 ± 16.63 ng/ml [gavage]; *p* <0.0001; **Figure 2**), indicating a lower stress response of the MDA glucose dosing method compared to gavage. There was no statistically significant sex effect **[Supplementary Table 1]** so data from both sexes were pooled.

**Figure 2:**
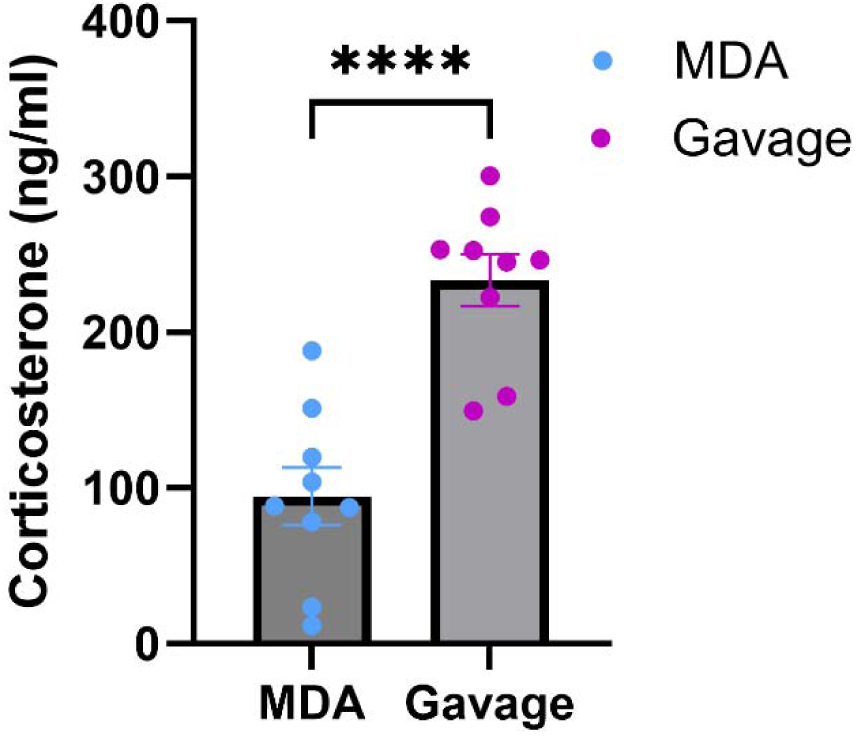
Lower plasma corticosterone after glucose-dosing by micropipette-guided drug administration (MDA) compared with oral gavage. Plasma corticosterone levels (ng/ml) in male and female C57BL/6J mice 10 minutes after dosing with 2.5g glucose/kg body weight of flavoured glucose solution. *p* <0.0001, unpaired t-test. N = 18 mice (11 male and 7 female). Data are presented as mean ± SEM, with circles indicating individual data points.

### MDA-glucose dosing produced a glucose clearance profile comparable to gavage-glucose dosing in an oGTT

Gavage is currently the standard method of dosing for a rodent oGTT. In mice, published data indicate that compared to the glucose clearance curve produced by an IPGTT, gavage dosing results in a lower glucose excursion in both lean and diet-induced obese mice, reflective of the incorporation of the incretin response to gavaged-glucose compared to IP-glucose^3^. These key differences highlight the importance of oral dosing in the GTT to assess the full physiological spectrum of glucose homeostasis, unless the specific goal of a study is to examine effects independent of the GI tract. We expected the MDA-oGTT to have a similar glucose clearance profile to the gavage-oGTT. Because of known differences in glucose homeostasis between C57BL6 sub-strains^15^, we compared oGTT profiles using MDA and gavage dosing in both C57BL/6J and C57BL/6N mice using a within-subjects randomised cross-over study.

Within sex and sub-strain, visually, the shape of the glucose clearance curves was broadly similar between MDA-oGTT and gavage-oGTT in both C57BL/6J and C57BL/6N mice [**Figure 3**]. There was a statistically significant effect of route of administration (RoA) in the female C57BL/6J mice (F_(1,5)_ = 6.7, *p*_(*RoA*)_ = 0.049), but not in male C57BL/6J mice (F_(1,7)_ = 3.2, *p*_(*RoA*)_ = 0.12) or either sex of C57BL/6N mice (males: F_(1,7)_ = 0.35, *p*_(*RoA*)_ = 0.57; females: F_(1,7)_ = 4.3, *p*_(*RoA*)_ = 0.078) [**Figure 3A, B, D and E**]. There was, as expected, an effect of time for all groups in both sub-strains (C57BL/6Js: males, F_(6, 42)_ = 48.1, *p*_(*time*)_ <0.0001; females, F_(2.9, 14.4)_ = 65.4, *p*_(*time*)_ <0.0001. C57BL/6Ns: males, F_(2.5, 17.6)_ = 28.7, *p*_(*time*)_ <0.0001; females, F_(6, 42)_ = 52.0, *p*_(*time*)_ <0.0001), but no interaction effect of time with sex or route of administration.

**Figure 3:**
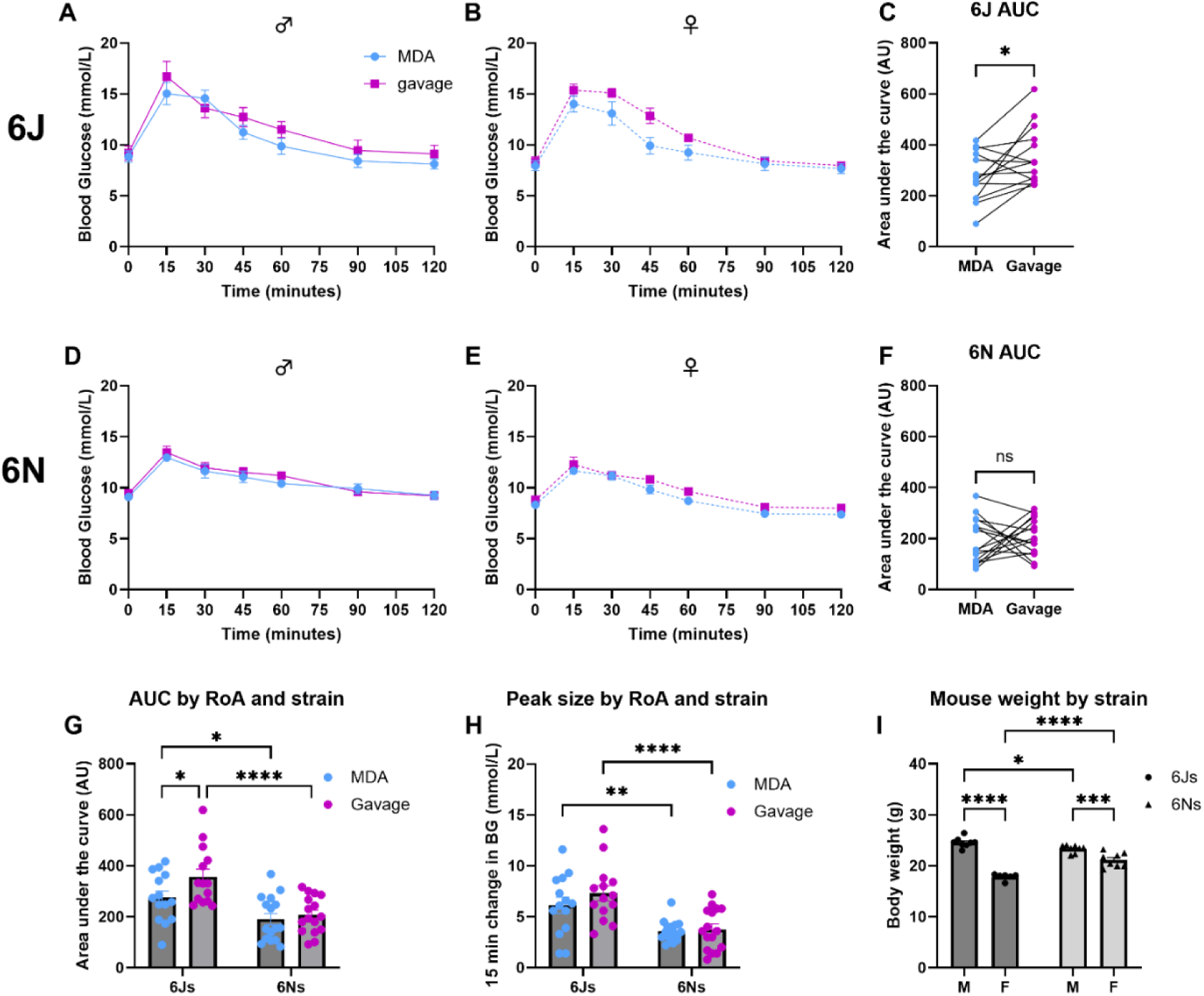
MDA-glucose dosing produced a glucose clearance profile comparable in shape to that of gavage-glucose dosing in an oral glucose tolerance test (oGTT) and was able to detect differences between C57BL/6J and C57BL/6N sub-strains. Using a within-subjects randomised cross-over design with order of route of administration balanced, male and female mice of both sub-strains were subjected to two oGTTs one-week apart. For each oGTT, animals were dosed orally with 2.5g glucose/kg body weight of flavoured glucose solution via gavage (magenta) or micropipette-guided drug administration (MDA; blue) after a basal blood glucose measurement (tail prick) at 0 minutes. Blood glucose was then measured at regular intervals for up to 120 minutes. For C57BL/6J mice (**A-C**) there was a statistically larger baseline subtracted area under the curve (AUC; **C**) in the gavage oGTT trial compared to the MDA-oGTT trial (\**p*<0.05, paired t-test). This was not observed in C57BL/6N mice (**D-F**). Comparisons between sub-strains were performed using a two-way repeated measures ANOVA for AUC (**G**) and peak glucose data (**H**), and a two-way ordinary ANOVA for body weight (**I**). For the AUC data (**G**) there were statistically significant effects of sub-strain (F _(1, 28)_ = 20.97, *p*_(*sub-strain*)_ <0.0001) and route of administration (RoA; F _(1, 28)_ = 4.66, *p*_(*RoA*)_ = 0.04), but no interaction between these factors. For the 15-minute peak glucose **(H)** there was a statistically significant effect of sub-strain (F _(1, 28)_ = 30.86, *p*_(*sub-strain*)_ <0.0001) but not RoA nor an interaction between these factors. For the body weight data **(I)** there was a statistically significant effect of sub-strain (F _(1, 26)_ = 7.70, *p*_(*sub-strain*)_ = 0.01), sex (F _(1, 26)_ = 146.0, *p*_(*sex*)_ <0.0001) and an interaction between the two variables (F _(1, 26)_ = 38.48, *p*_(*sub-strain*_ *_x sex_*_)_ <0.0001). **G-I**, post-hoc comparisons: * *p*<0.05; ** *p*<0.01; *** *p*<0.001; **** *p*<0.0001. n.s. Not statistically significant. N = 14 C57BL/6J mice (8 male and 6 female) and N = 16 C57BL/6N mice (8 male and 8 female). Data are presented as mean ± SEM with circles indicating individual data points.

Interestingly, in C57BL/6J mice there was a significant difference in the baseline subtracted area under the curve (AUC) for the glucose clearance profile during the gavage-oGTT compared to the MDA-oGTT, where gavage-oGTT produced a significantly greater AUC when analysed by a paired t-test where each mouse was compared to themselves after receiving both dosing methods, and by two-way repeated measures ANOVA [**Figure 3C**; F_(1,12)_ = 6.29, *p_(RoA)_* = 0.028]. Using the same analysis approaches, there was no statistically significant difference in the baseline subtracted oGTT AUCs between routes of administration for C57BL/6N mice [**Figure 3F**; F_(1,14)_ = 0.25, *p_(RoA)_* = 0.62]. There was also no effect of sex on the AUC for either sub-strain.

When comparing between sub-strains, there was no difference in basal blood glucose, however C57BL/6J mice showed a statistically greater baseline subtracted AUC (F_(1,28)_ = 21.0, *p_(sub-strain)_* <0.0001) and peak glucose excursion (sub-strain: F_(1,28)_ = 30.9, *p_(sub- strain)_* <0.0001) than C57BL/6N mice [**Figure 3G, H; Supplementary Table 2]**. There were no statistically significant interactions between sub-strain and route of administration on either baseline subtracted AUC or peak glucose excursion.

At 9-weeks of age (timing of first oGTT), there was a statistically significant effect of sub-strain (*F*_(1,26)_ = 7.69, *p_(sub-strain)_ =* 0.01) and sex (*F*_(1,26)_ = 145.9, *p_(sex)_* <0.0001) on body weight between the C57BL/6J and C57BL/6N mice used in this experiment [**Figure 3I; Supplementary Table 2]** and a significant interaction between the variables (*F*_(1,26)_ = 38.5 *p_(sub-strain x sex)_* <0.0001).

Here we have shown that MDA is a valid method of supplying a glucose bolus dose for the oGTT, which produces a reliable peak and glucose clearance curve and is able to replicate differences in glucose homeostasis reported between C57BL/6 sub-strains using other glucose dosing methods^15–18^.

### MDA-oGTT detected a change in GTT profile in high-fat diet-fed (HFD) mice

We tested whether we could detect impaired glucose tolerance by performing oGTTs in HFD-fed mice using MDA dosing. We performed a within-subjects randomised crossover oGTT study in male and female C57BL/6J mice to compare glucose clearance curves in response to MDA-glucose and gavage-glucose dosing, before and after a 16-week high-fat diet [**Figure 4A-F**].

**Figure 4:**
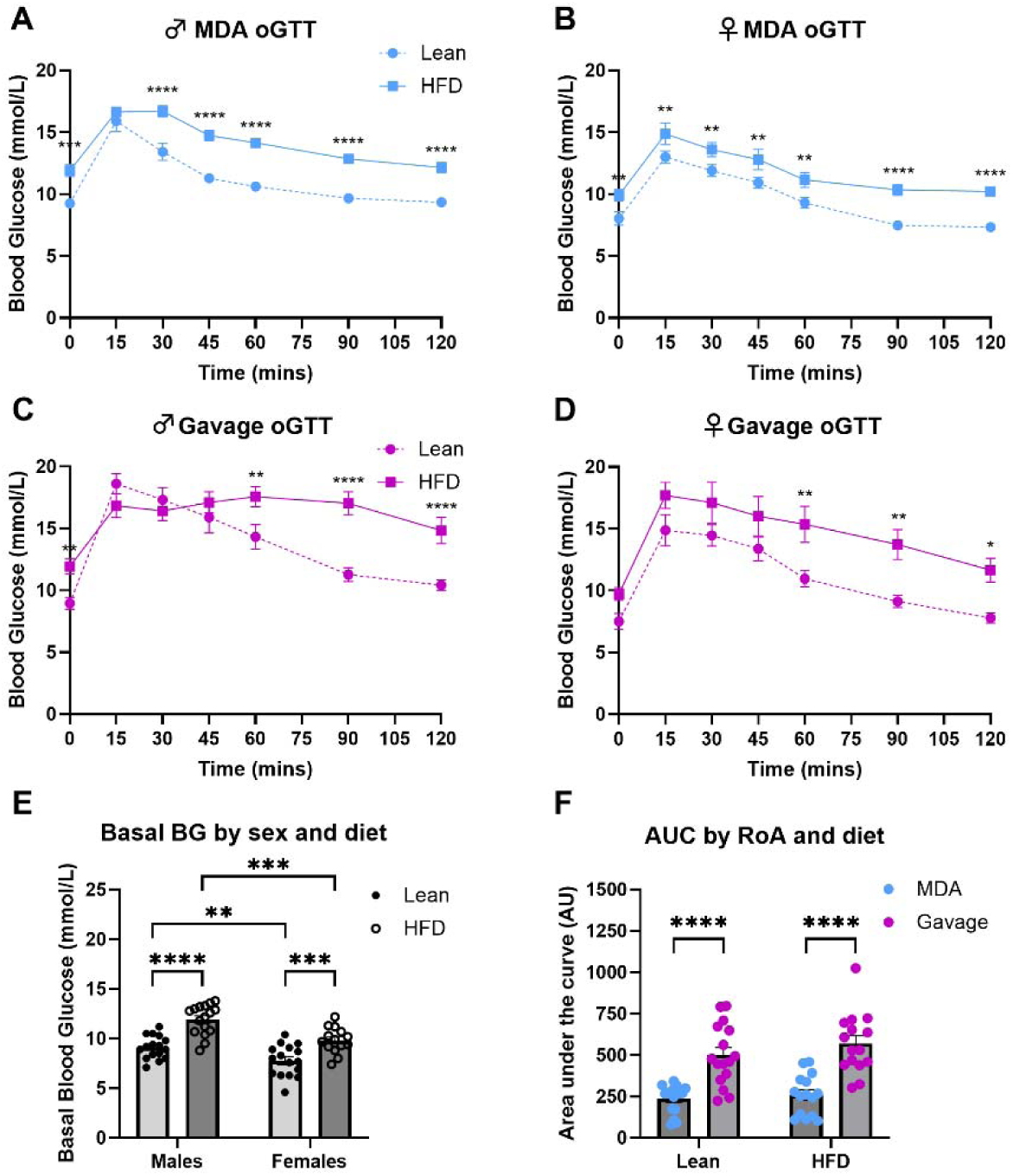
MDA-glucose revealed differences in glucose clearance in an oral glucose tolerance test (oGTT) in C57BL/6J mice fed a high-fat diet (HFD) for 16-weeks. Using a within-subjects randomised cross-over design with order of route of administration balanced, male and female C57BL/6J mice were subjected to two oGTTs one-week apart before and after receiving a HFD as the only source of food for 16-weeks. For each oGTT, animals were dosed orally with 2.5g glucose/kg body weight of flavoured glucose solution via gavage (magenta) or micropipette-guided drug administration (MDA; blue) after a basal blood glucose measurement (tail prick) at 0 minutes. Blood glucose was then measured regularly for 120 minutes. Comparisons of glucose clearance curves within sex **[A-D]** were performed using a two-way repeated measures ANOVA with Sidaks-test *post-hoc* analysis applied. MDA-oGTT detected an effect of diet on glucose clearance profile in male [**A**; F_(1, 7)_ = 61.36, *p*_(*diet*)_ = 0.0001] and female mice [**B**; F_(1, 6)_ = 17.15, *p*_(*diet*)_ = 0.0061]. Gavage-oGTT did not reveal statistically significant changes in glucose clearance in male mice (**C**; F_(1, 7)_ = 5.007, *p*_(*diet*)_ = 0.0603) but did in females [**D**; F_(1, 7)_ = 16.98, *p*_(*diet*)_ = 0.0045]. Basal blood glucose was significantly elevated in the oGTT after the 16-week HFD **[E]** (F_(1, 58)_ = 46.59, *p*_(*diet*)_ <0.0001), and showed a significant sex effect (F_(1, 58)_ = 22.98, *p*_(*sex*)_ <0.0001). After baseline subtraction to adjust for differences in basal blood glucose, there was a statistically significant effect of route of administration (F _(1, 15)_ = 64.70, *p_(RoA)_* <0.0001) but not diet (F _(1, 15)_ = 1.58, *p_(diet)_* = 0.23) on area under the curve (AUCs) between lean and HFD-fed C57BL/6J mice **[F]**. There was no statistical interaction between the variables (F _(1, 13)_ = 0.44, *p_(RoA x diet)_* = 0.52). N = 15 (8 male and 7 female) mice. Data are presented as mean ± SEM, with circles indicating individual data points.

In the lean C57BL/6J animals (prior to switching to the HFD), when comparing MDA-oGTT and gavage-oGTT glucose clearance curves there was a statistically significant effect of route of administration (males; F_(1, 14)_ = 13.36, *p*_(*RoA*)_ = 0.0026 and females; F_(1, 14)_ = 5.96, *p*_(*RoA*)_ = 0.029), time (males; F_(2.7, 38.25)_ = 64.72, *p*_(*time*)_ <0.0001 and females; F_(2.7, 37.8)_ = 82.9, *p*_(*time*)_ <0.0001), and an interaction between route of administration and time (males; F_(6, 84)_ = 5.52, *p*_(*RoA*_ *_x time_*_)_ <0.0001 and females; F_(6, 84)_ = 3.54, *p*_(*RoA*_ *_x time_*_)_ = 0.0036) **[Supplementary Figure 2A and B**]. Baseline subtracted AUC showed a significant effect of route of administration (F_(1, 14)_ = 31.93, *p*_(*RoA*)_ <0.0001) but not sex, nor an interaction between these two variables **[Supplementary Figure 2C**]. Together, these data broadly reproduce those of the first study in C57BL/6J mice [**Figure 3**].

On the HFD, the mean body mass increase of males was 79.35 ± 5.42 % (Change: 27.56 ± 0.74 g to 49.43 ± 1.88 g), and females gained 94.82 ± 6.20 % (Change: 20.50 ± 0.34 g to 40.03 ± 1.81 g). Analysis of the body weight data revealed a statistically significant effect of diet (F_(1, 13)_ = 370.6, *p*_(*diet*)_ <0.0001) and sex (F_(1, 13)_ = 25.3, *p*_(*sex*)_ = 0.0002), but there was no significant interaction between the two variables.

Feeding a HFD induced changes in glucose clearance profiles in C57BL/6J mice. Both MDA-oGTT (F_(1, 6)_ = 17.15, *p*_(*diet*)_ = 0.0061) and gavage-oGTT (F_(1, 7)_ = 16.98, *p*_(*diet*)_ = 0.0045) detected statistically significant effects of diet on glucose clearance profiles in female mice. Only MDA detected a statistically significant effect of diet in the male mice (F_(1, 7)_ = 61.36, *p*_(*diet*)_ = 0.0001). However, there were interesting differences in the shape of the glucose tolerance curve profiles of the HFD-fed mice between sexes, where HFD-fed males tended to have a prolonged peak in the MDA profile, and delayed clearance in the gavage profile not seen in HFD-fed female animals [**Figures 4A-D**]. These differences are reflected by statistically significant interactions between diet and time in male mice when receiving both MDA-oGTT (F_(6, 42)_ = 2.97, *p*_(*time*_ *_x diet_*_)_ = 0.0165) and gavage-oGTT (F_(6, 42)_ = 10.19, *p*_(*time*_ *_x diet_*_)_ <0.0001), that were not observed in the female animals. As expected, there was a statistically significant effect of time in all groups (male MDA-oGTT: F_(6, 42)_ = 107.7, *p*_(*time*)_ <0.0001; male gavage-oGTT: F_(6, 42)_ = 39.48, *p*_(*time*)_ <0.0001; female MDA-oGTT: F_(6, 36)_ = 59.21, *p*_(*time*)_ <0.0001; female gavage-oGTT: F_(6, 42)_ = 36.17, *p*_(*time*)_ <0.0001).

Compared to the lean state, both male and female C57BL/6J mice had significantly elevated basal blood glucose in the oGTT after the 16-week HFD [**Figure 4E** (F_(1, 58)_ = 46.59, *p*_(*diet*)_ <0.0001)], and there was a significant sex effect (F_(1, 58)_ = 22.98, *p*_(*sex*)_ <0.0001), but no statistical interaction between sex and diet. After baseline subtraction to adjust for differences in basal blood glucose, there was an effect of route of administration (F _(1, 15)_ = 64.70, *p_(RoA)_* <0.0001) but not diet (F _(1, 15)_ = 1.58, *p_(diet)_* = 0.23) on AUCs between lean and HFD-fed C57BL/6J mice [**Figure 4F**]. There was no statistical interaction between the variables (F _(1, 13)_ = 0.44, *p_(RoA x diet)_* = 0.52).

### Inter-institutional reproducibility of the MDA-oGTT assay

A key aspect of developing a new technique is to ensure that it can be reproduced in separate institutions with the same result. To investigate this, at Vanderbilt MMPC-Live we carried out a within-subjects randomised crossover in C57BL/6J mice as described above, with a few minor differences which reflected the standard practice locally: the method of blood sampling and the number of handlers (see methods section for details). As at Exeter, at Vanderbilt MMPC-Live all mice readily habituated to MDA-glucose and drank with either a gentle tail hold or unrestrained on the day of the oGTT. Within sex, MDA-oGTT produced a peak blood glucose at 15 minutes and had a similar profile to the gavage-oGTT [**Figure 5A and B**]. There was a statistically significant effect of route of administration in females (males: F _(1, 7)_ = 4.62, *p_(RoA)_* = 0.068; females: F _(1, 7)_ = 16.94, *p_(RoA)_* = 0.0045). Again, we showed that gavage-oGTT generated a significantly larger baseline subtracted AUC than MDA-oGTT when mice were compared to themselves through a paired t-test [**Figure 5C**; *p* = 0.0015]. However, within a two-way repeated measures ANOVA of the individual data sets only females showed a significant difference between the AUCs of gavage- and MDA-oGTT [**Figure 5D**].

**Figure 5:**
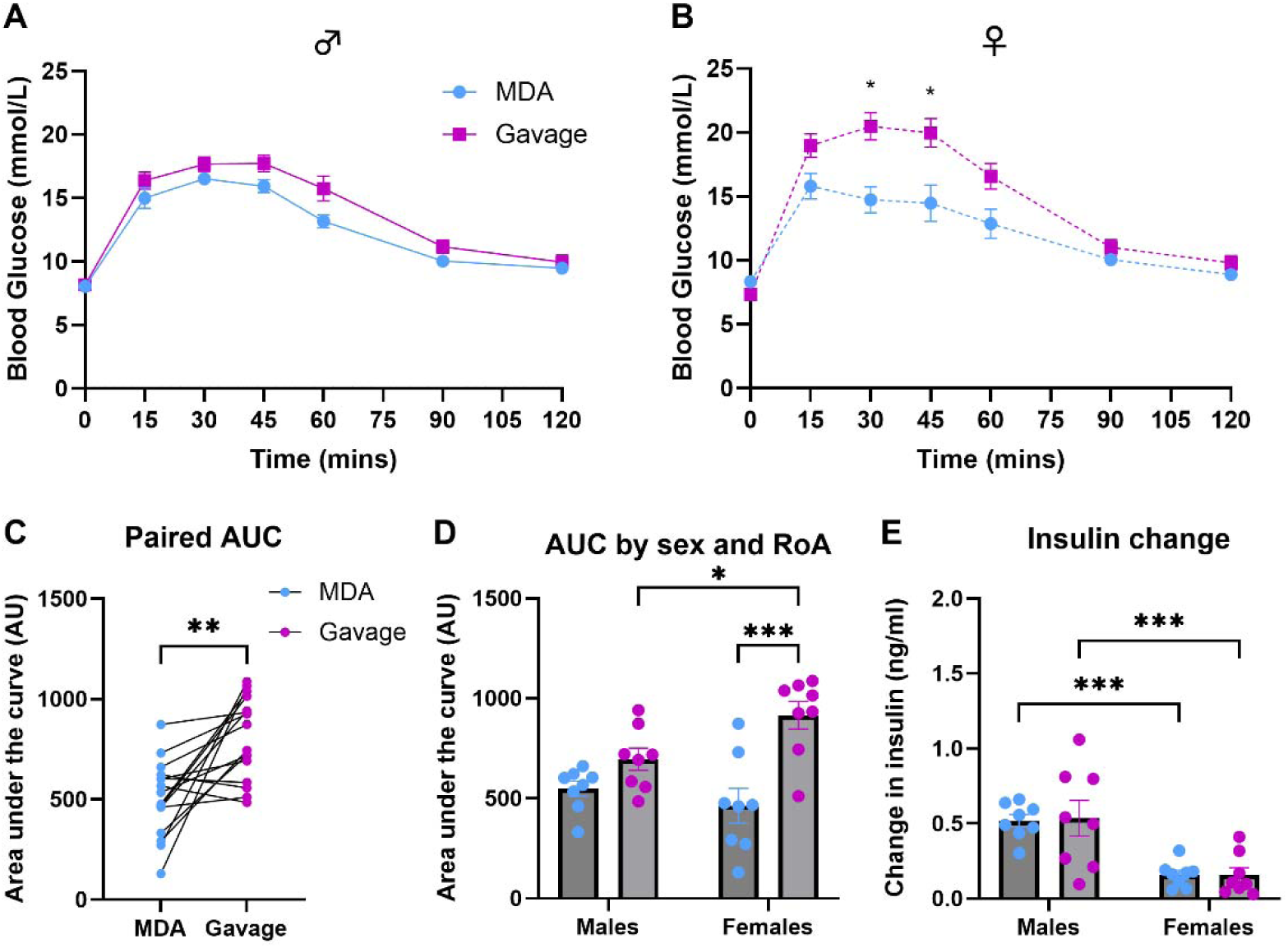
MDA-glucose dosing produced a glucose clearance profile comparable in shape to that of gavage-glucose dosing in an oral glucose tolerance test (oGTT) in C57BL/6J mice in studies performed at Vanderbilt MMPC-Live. Using a within-subjects randomised cross-over design with order of route of administration balanced, male and female C57BL/6J were subjected to two oGTTs one-week apart. For each oGTT, animals were dosed orally with 2.5g glucose/kg body weight of flavoured glucose solution via gavage (magenta) or micropipette-guided drug administration (MDA; blue) after a basal blood glucose measurement (tail-tip excision) at 0 minutes. Blood glucose was then measured at regular intervals for up to 120 minutes. Within-sex analysis of glucose clearance curves using a two-way repeated measures ANOVA indicated a statistically significant effect of route of administration (RoA) in females but not males (**[A]** males: F _(1, 7)_ = 4.62, *p_(RoA)_* = 0.068; **[B]** females: F _(1, 7)_ = 16.9, *p_(RoA)_* = 0.0045). Gavage-oGTT generated a significantly larger baseline subtracted AUC than MDA-oGTT when mice were compared to themselves through a paired t-test, \*\**p* = 0.0015 **[C]**. The sex-separated AUC data **[D]** were analysed by two-way repeated measures ANOVA for sex and route of administration effects, which indicated a significant effect of route of administration (F _(1, 14)_ = 18.9, *p_(RoA)_* = 0.0007) as well as an interaction between route of administration and sex (F _(1, 14)_ = 4.83, *p_(interaction)_* = 0.045). The change in plasma insulins between the samples taken 15 minutes before and after the glucose bolus was calculated for each mouse **[E]** and there was a significant effect of sex (F _(1, 14)_ = 24.04, *p_(RoA)_* = 0.0002) but not route of administration (F _(1, 14)_ = 0.013, *p_(RoA)_* = 0.91). **D and E** post-hoc comparisons: * *p*<0.05; ** *p*<0.01; *** *p*<0.001. N = 16 C57BL/6J mice (8 male and 8 female). Data are presented as mean ± SEM.

At Vanderbilt MMPC-Live larger blood samples were collected to enable assessment of plasma insulin. The change in plasma insulins between the samples taken 15 minutes before [MDA, 0.34 ± 0.01 ng/ml (males) and 0.28 ± 0.02 ng/ml (females); Gavage, 0.46 ± 0.04 ng/ml (males) and 0.26 ± 0.03 (females)] and after the glucose bolus [MDA, 0.86 ± 0.05 ng/ml (males) and 0.43 ± 0.04 ng/ml (females); Gavage, 1.00 ± 0.14 ng/ml (males) and 0.42 ± 0.07 (females)] were calculated for each mouse. There was a significant effect of sex (F _(1, 14)_ = 24.04, *p_(sex)_* = 0.0002) but not route of administration (F _(1, 14)_ = 0.013, *p_(RoA)_* = 0.91) on the change in plasma insulin [**Figure 5E**]. There was no statistically significant interaction between the variables.

## Discussion

Here, we have validated MDA as a viable alternative oral glucose dosing method for the oGTT to that of the more invasive gavage. Although some potentially interesting differences were observed, we have broadly shown that MDA-glucose administration produced a comparable glucose clearance profile to gavage-glucose in the oGTT in lean C57BL/6J and C57BL/6N mice, and we were able to replicate published sub-strain differences in glucose tolerance^15–18^. Obesity-associated differences in oGTT profiles were observed using both MDA-glucose and gavage-glucose, although a longer HFD-feeding period and/or use of a different diet may have been beneficial to enable more pronounced differences to become evident. Critically, we have shown significantly lower plasma corticosterone levels in the MDA-glucose treated mice compared to those dosed via gavage, indicating a reduction in procedural stress, which may contribute to lesser inter-animal variation within the glucose measurements of the MDA-treated mice. Together these data support the MDA-oGTT method as a refinement in terms of both animal experience and data quality. Due to lower experimental variation, use of this method may allow for a reduction in the number of mice required for this type of metabolic study and/or enable more subtle physiological effects to be observed.

When compared to intraperitoneal injection, oral gavage dosing of glucose for a GTT is more physiologically relevant; however, it is invasive, causes stress, and can be difficult for inexperienced investigators to carry out correctly. Gavage dosing also fails to incorporate a potentially crucial part of the body’s homeostatic response to food – that of the cephalic stage. Cephalic-phase responses are activated by the detection of an oral food stimulus by chewing/licking, taste, smell and even sight, and facilitate preparation for the metabolism of substances prior to and during ingestion^4^. These responses contribute to the complex web of metabolic signalling in numerous ways and are sustained throughout a meal^4^. An oral gavage, where the glucose is deposited directly into the stomach whilst bypassing taste and olfactory receptors, would not initiate the full range of cephalic phase responses to a physiological degree, and therefore responses during a metabolic study, such as a GTT, are likely different relative to the voluntary oral ingestion of the stimulus. The MDA method however, where mice see, smell, lick, taste, and willingly drink the glucose bolus, incorporates the full range of cephalic-phase responses, and therefore any metabolic data gained from MDA-dosed mice should be more physiologically accurate and translationally relevant to the human clinical assay (where people voluntarily drink a sweetened drink) than data obtained from an oral gavage.

The C57BL/6N and C57BL/6J sub-strains, both commonly used across *in vivo* metabolism research, share an ancestral background but differ in regard to the *Nnt* gene – where 6Ns carry the wild-type gene, and 6Js carry a mutated version with a 5-exon deletion as well as a missense mutation^19^. *Nnt* encodes nicotinamide nucleotide transhydrogenase (NNT), an inner mitochondrial membrane protein involved in the catalysis of NADPH production and proton translocation across the inner mitochondrial membrane^20^. Ultimately, the mutation is associated with impaired glucose tolerance and a reduction in glucose-stimulated insulin secretion in C57BL/6J mice^15^, which contributes to reported differences in the GTT profiles of the sub-strains^15–18^. In our studies in C57BL/6J mice, integration of insulin-independent cephalic-phase responses may be an explanation for the observed differences in glucose clearance profiles in response to the MDA-oGTT (in both the lean and obese state) compared with gavage-oGTT; however, the physiological mechanism(s) underlying these differences remain to be explored in future studies.

As we chose to dose glucose to body weight, it is possible that the significant difference in body weights at 9-weeks of age is a contributing factor to the differences in GTT profiles seen between C57BL/6N and C57BL/6J sub-strains. However, this is less likely to be impactful when comparing between lean animals than when comparing lean and obese animals or other models where body composition, in particular the balance between fat and lean mass, is known to be altered^21^. While it should be noted that this sub-strain comparison was not performed on animals who received oGTT during the same experiment (i.e., GTTs for each strain were carried out on different days, albeit with the same handler, environment, and time of day), the observed sub-strain differences are in-line those reported in the literature where direct within-experiment sub-strain comparisons were performed^15–18^. Importantly, in the context of this study, within sub-strain, the oGTT profiles for MDA-dosed and gavage-dosed C57BL/6N mice are comparable to those reported in the literature^3^.

There are other published experimental approaches which have demonstrated refinements of oGTT protocols to include voluntary glucose consumption. These include timed-ingestion of flavoured-sweetened gels^13^ and voluntary licking of a glucose solution after water-deprivation^22^. A challenge of the gel-consumption approach is variability in the speed and consistency of gel-consumption within the desired period ahead of the GTT, which can lead to some animals being excluded on procedural grounds (Kennard *et al*^13^, and *A. King personal communication*). During our study, of the 107 animals that were given MDA-glucose, none failed to consume the full glucose dose on the day of the oGTT. Although similar in some ways, our study has advantages over the approach of Glendinning *et al* that trained animals to lick glucose from a sipper, as it does not require water deprivation^22^, which may cause stress and impact other related physiological processes.

Finally, we have shown that the MDA technique can be utilised for an oGTT in a separate institution, demonstrating cross-institutional reproducibility of the method. Interestingly, the glucose clearance profiles of both the oGTTs for both MDA and gavage at Vanderbilt MMPC-Live showed a prolonged peak, which was not seen in the data generated at the University of Exeter but was likely a result of the different method of blood sampling used. Where the blood samples taken in **Figures 3 and 4** were from a small needle prick in the tip of the tail, those in **Figure 5** were taken after excision of the tip of the tail, which enabled the larger blood sampling required for measuring plasma insulins. Despite local anaesthetic cream being used on the tail, this was undoubtedly a stressor for the mice involved and evidence of how changing certain parts of an oGTT method can affect the profile of the data.

Of course, any change in experimental protocols brings new points of discussion. In this case, one area for reflection is the required mouse habituation time. Of note here, we used the tunnel handling/hand cupping method during habituation and husbandry for our study which, compared to traditional tail-handling, is shown to reduce aversion and anxiety associated with investigator interactions^10–12^. While it is already strongly recommended for researchers to habituate rodents to handling ahead of a GTT to reduce procedural/experimental stress, the MDA-oGTT method potentially requires some additional period of habituation time, though in our experience this should not necessarily be more than one week. This means an increased requirement for researcher time, which may be seen as a limitation; however, our experience is that the researcher time required during the handling and dosing-habituation weeks is typically <2 mins per animal per day. The trade-off of the additional time should be weighed carefully against the benefits of the refinement, encompassing greater translatability to the human clinical assay, more holistic physiological relevance, and improved animal welfare and investigator experience. In an era of animal research where we are striving for continuous refinement of study design, an increase in researcher time should not be seen as a block but rather part of the normal process for securing the highest quality data and welfare standards.

There is a risk with any habituation process of physiological conditioning, which leads to the animals associating the appearance of the investigator with receiving glucose, potentially leading to anticipatory responses such as insulin secretion. At this time, we have no evidence to support or refute this possibility; however, it could be argued that this may happen in the human clinical setting as the patient knows before going to the clinic that they will be drinking a sweet drink as part of the GTT. Future studies exploring how long the memory of habituation to MDA-glucose is retained may enable increased gaps between habituation and testing that could minimise concerns relating to conditioning. It is our view that the benefits of the MDA-oGTT (outlined above) outweigh these concerns.

In summary, we have developed a more refined method for the oGTT using MDA-glucose delivery that has greater clinical translatability and has benefits for both animal and investigator experience. Additionally, lower inter-animal variability observed using this method may enable fewer animals to be used per study and/or smaller effect sizes to be observed. This simple approach requires no specialist equipment and minimal training, so should be readily adoptable by others.

## Acknowledgements

The authors are keen to share detailed protocols and their experience with others to promote widespread adoption of the method, so please do not hesitate to get in contact with us by email. All data are available on reasonable request.

The authors would like to thank the staff at the University of Exeter BRF for their support and care of the animals. We are also grateful to Prof Aileen King and Dr Chloe Rackham for helpful discussions and support relating to blood sampling, and the technical staff of the Vanderbilt Mouse Metabolic Phenotyping Center-Live for their help generating some of the data herein. This project was supported by a grant from the National Centre for Replacement, Reduction and Refinement of the use of animals in Research (NC3Rs; NC/X000923/1; to KE and CB) and Diabetes UK (19/0006035; to KE and CB). The MMPC at Vanderbilt is supported by DK135073 (MMPC-Live) and DK020593 (DRTC).

This paper is dedicated to the memory of Prof David Wasserman of Vanderbilt University School of Medicine, a leading figure in setting standards for mouse metabolic phenotyping and understanding the physiological basis of glucose homeostasis, but also a great friend and mentor.

## Supplementary Figures

**Supplementary Figure 1:**
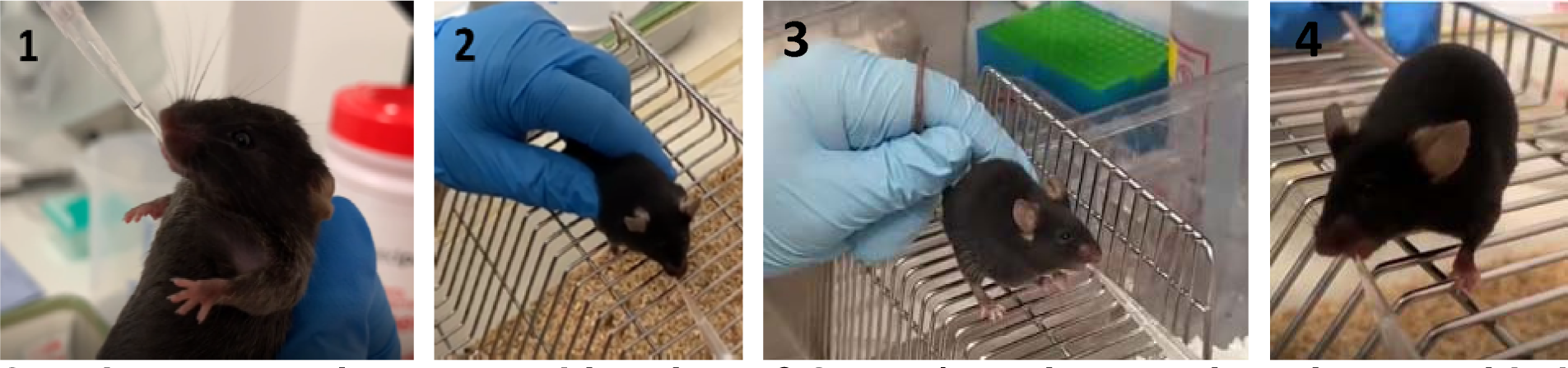
Habituation of C57BL/6J mice to micropipette-guided dosing with chocolate-flavoured glucose (40% solution with flavouring) **1**: Mice were restrained via scruff and the pipette containing the glucose solution was presented at a horizontal angle. **2**: Mice were gently restrained atop the hopper with a light grip on their tail and above their hind legs, and the pipette was once again presented at a horizontal angle. **3**: Mice were held gently by the tail and the pipette was presented as before. **4**: Mice were placed atop the hopper and drank from the pipette without restraint.

**Supplementary figure 2:**
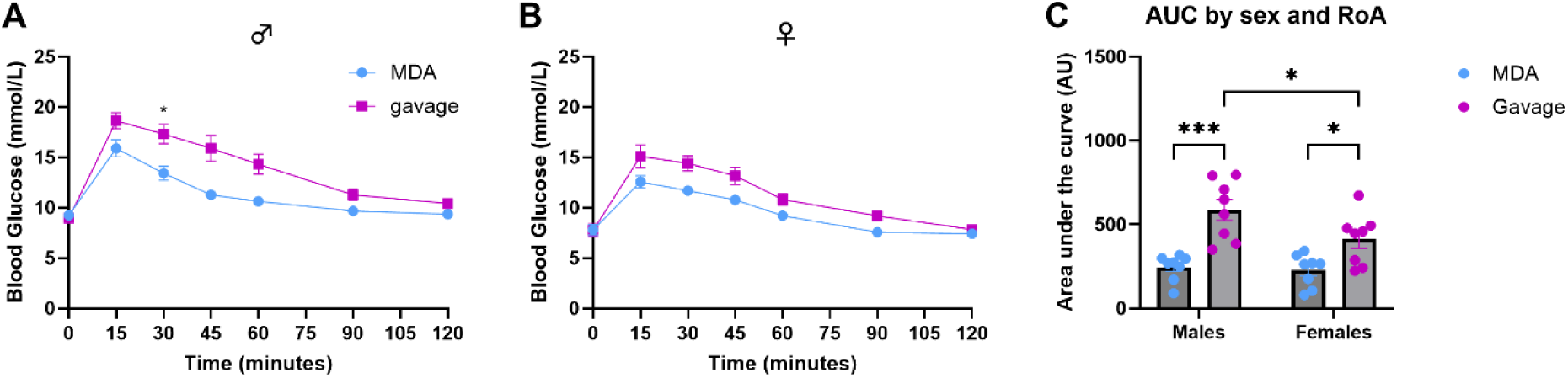
MDA-glucose dosing produced a glucose clearance profile comparable in shape to that of gavage-glucose dosing in an oral glucose tolerance test (oGTT) in lean C57BL/6J mice (prior to introduction of the high-fat diet) Using a within-subjects randomised cross-over design with order of route of administration balanced, male and female C67BL/6J mice (8 males, 8 females) were subjected to two oGTTs one-week apart prior to the introduction of a high-fat diet (see **figure 4**). For each oGTT, animals were dosed orally with 2.5g glucose/kg body weight of flavoured glucose solution via gavage (magenta) or micropipette-guided drug administration (MDA; blue) after a basal blood glucose measurement (tail prick) at 0 minutes. Blood glucose was then measured at regular intervals for up to 120 minutes. For both sexes (**A-C**) there was a statistically larger baseline subtracted area under the curve (AUC; **C**) in the gavage-oGTT compared to the MDA-oGTT, and a sex difference in the AUC of the gavage, but not the MDA-oGTT. AUC data analysed by two-way repeated measures ANOVA for effects of sex (F _(1, 14)_ = 4.13, *p_(sex)_* = 0.06), and route of administration (RoA; F _(1, 14)_ = 31.9, *p_(RoA)_* <0.0001), with multiple comparisons (* *p*<0.05; *** *p*<0.001). Data are presented as mean ± SEM.

## Supplementary Tables

**Supplementary table 1:** Effect of sex and route of administration (RoA) on plasma corticosterone. Data analysed by 2-way repeated measures analysis of variance (RM-ANOVA) for sex, RoA interaction. Bolded values indicate statistical significance (*p*<0.05).

| Figure 2 | N | Sex | RoA | RoA x Sex |
| --- | --- | --- | --- | --- |
| C57BL/6Js | 18<br>(11m, 7f) | $F_{(1,14)} = 1.98$<br>$p = 0.18$ (ns) | **$F_{(1,14)} = 30.9$**<br>**$p < 0.0001$ (\*\*\*\*)** | $F_{(1,14)} = 0.014$<br>$p = 0.91$ (ns) |

**Supplementary table 2:**
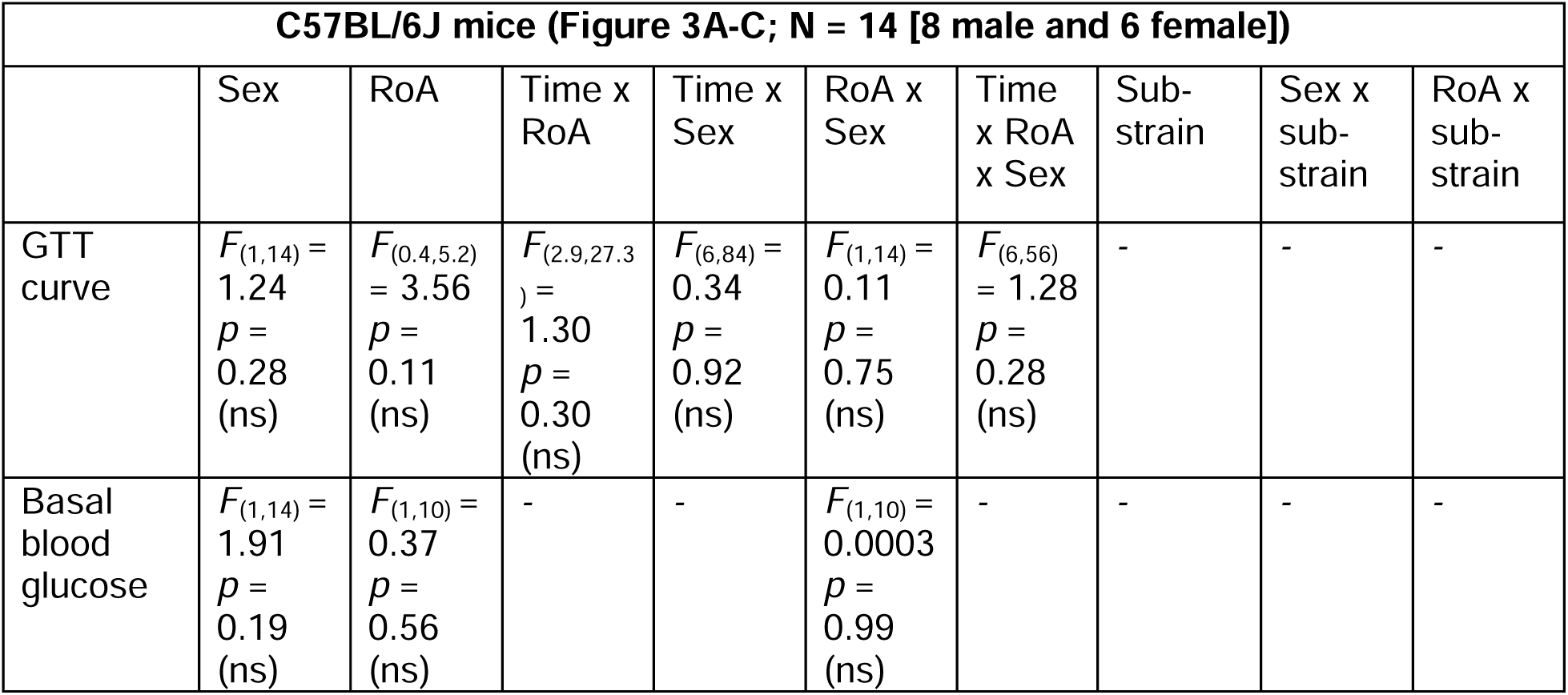
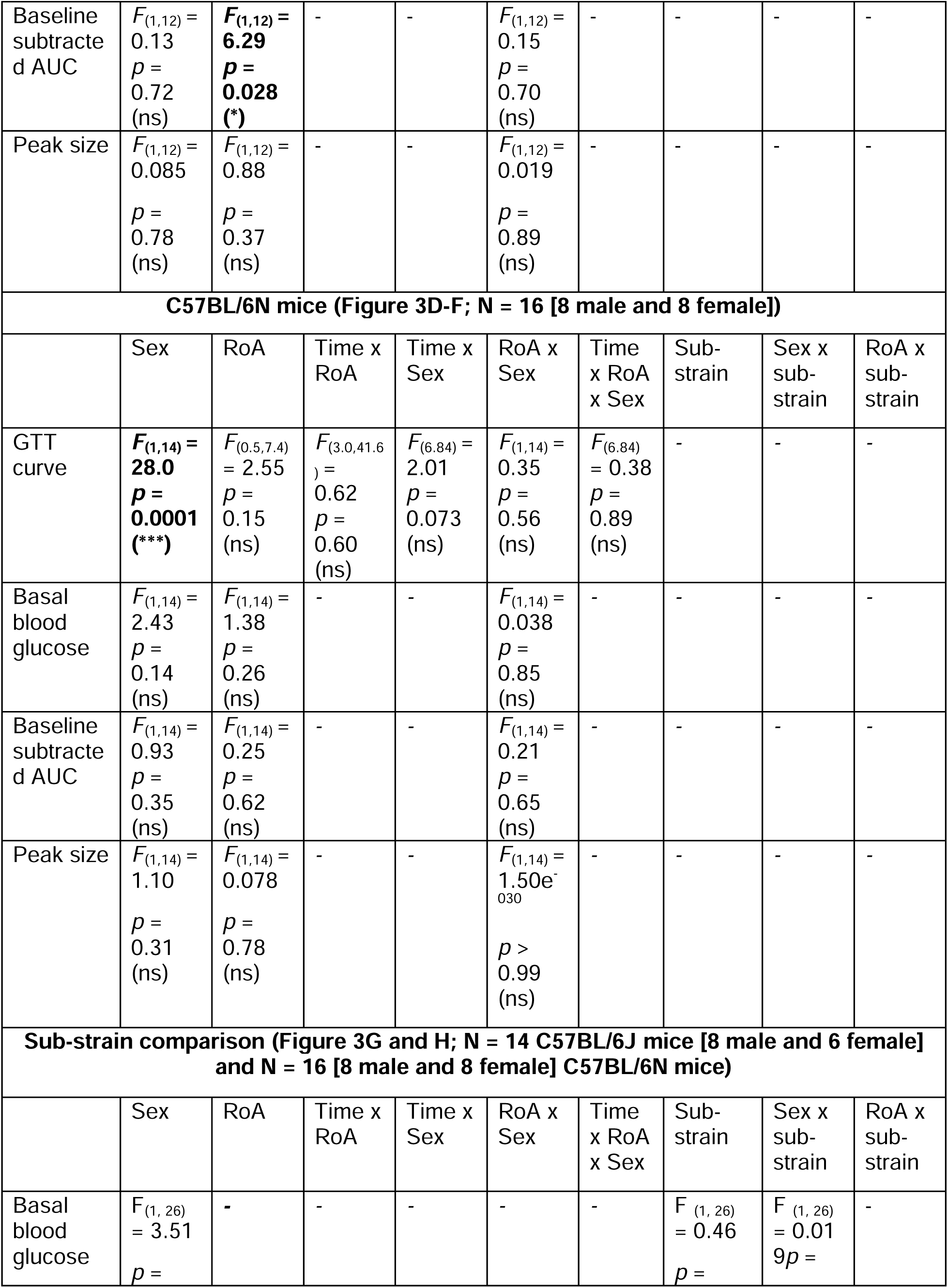
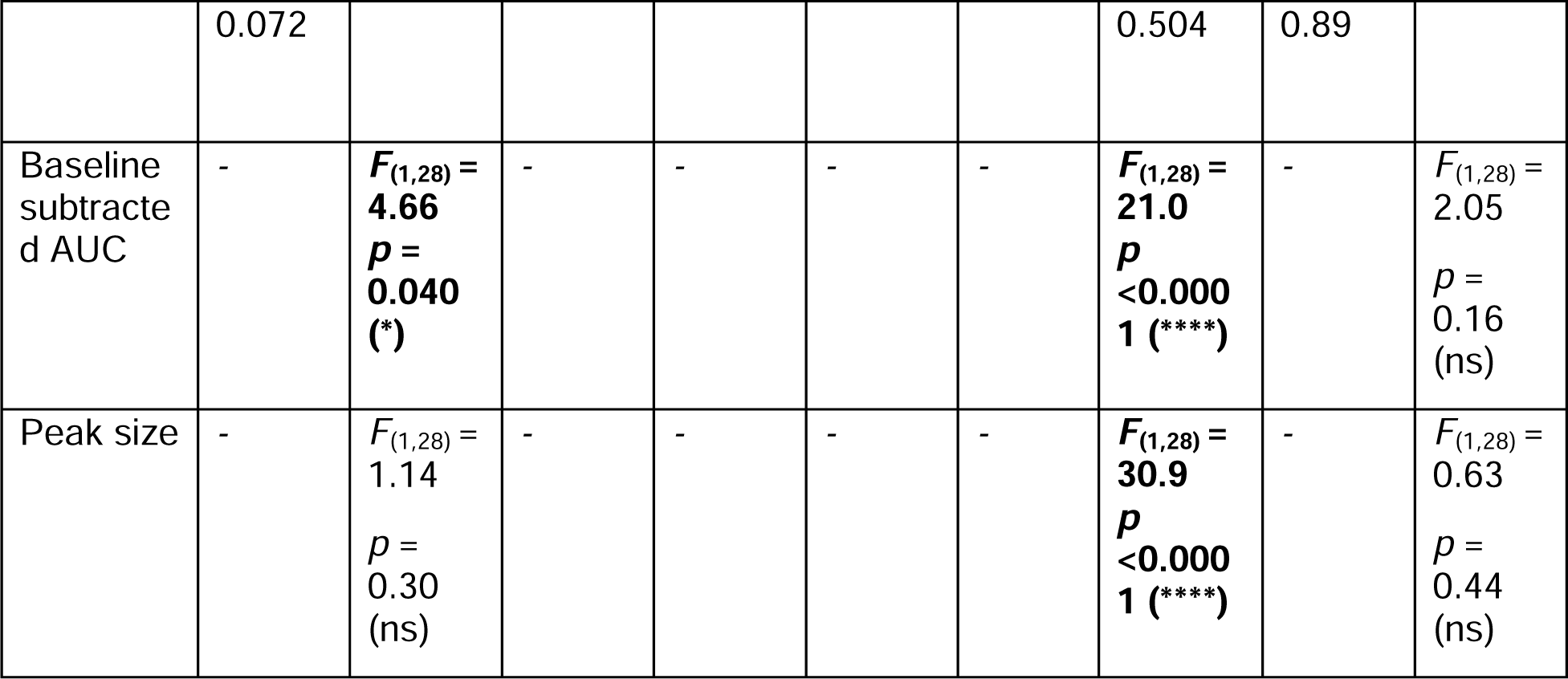
Comparisons of C57BL/6J and C57BL/6N data. Glucose tolerance test (GTT) curve data for each strain after a randomised crossover, where each mouse was given both a MDA-oGTT and a gavage-oGTT one week apart, with the order of dosing routes randomised. Data were analysed by 3-way repeated measures analysis of variance (RM-ANOVA) for time, sex, and route of administration (RoA) interaction with Tukey’s multiple comparisons test. Area under curve (AUC), basal blood glucose (BG), and peak size data for each strain were analysed by 2-way RM-ANOVA for sex and RoA interaction. AUC and peak size strain comparisons were analysed by 2-way RM-ANOVA for RoA, strain interaction. Bolded values indicate statistical significance (*p*<0.05). ns = not statistically significant (*p*>0.05).

**Supplementary table 3a:** HFD data. C57BL/6J mice (n= 8 males, 7 females) were given two oGTTs (MDA-oGTT and gavage-oGTT one week apart) in a randomised crossover before and after a 16-week high fat diet (HFD). Lean and HFD GTT data were analysed by 3-way repeated measures analysis of variance (RM-ANOVA) for time, sex, route of administration (RoA) interaction with Tukey’s multiple comparison test. Comparisons of area under curve (AUC), basal blood glucose (BG) and peak size between diets were analysed by 3-way RM-ANOVA for sex, RoA, diet interaction with Tukey’s multiple comparisons test. Bolded values indicate statistical significance (*p*<0.05).

| C57BL/6J mice (Figure 4A-F; N = 15 [8 male and 7 female]) |  |  |  |  |  |  |  |  |  |  |
| --- | --- | --- | --- | --- | --- | --- | --- | --- | --- | --- |
|  | Sex | RoA | Diet | Time x RoA | Time x Sex | RoA x Sex | RoA x Diet | Diet x sex | Time x RoA x Sex | Diet x RoA x Sex |
| Lean GTT curve | $F_{(1,14)} = 15.1$<br>$p = 0.0017$<br>(**) | $F_{(0.5,7.2)} = 34.2$<br>$p = 0.0015$<br>(**) | - | $F_{(2.7,37.8)} = 7.22$<br>$p = 0.0009$<br>(***) | $F_{(6,84)} = 2.46$<br>$p = 0.031$<br>(*) | $F_{(1,14)} = 1.54$<br>$p = 0.24$<br>(ns) | - | - | $F_{(6,84)} = 0.98$<br>$p = 0.44$<br>(ns) | - |
| HFD GTT curve | $F_{(1,13)} = 8.92$<br>$p = 0.011$<br>(*) | $F_{(0.5,6.6)} = 10.6$<br>$p = 0.024$<br>(*) | - | $F_{(3.6,46.3)} = 7.81$<br>$p = 0.0001$<br>(***) | $F_{(6,78)} = 2.09$<br>$p = 0.064$<br>(ns) | $F_{(1,13)} = 0.37$<br>$p = 0.56$<br>(ns) | - | - | $F_{(6,78)} = 3.41$<br>$p = 0.0049$<br>(**) | - |
| Basal subtracted AUCs | $F_{(1,14)} = 0.92$<br>$p = 0.35$<br>(ns) | $F_{(1,14)} = 68.1$<br>$p < 0.0001$<br>(****) | $F_{(1,14)} = 1.82$<br>$p = 0.20$<br>(ns) | - | - | $F_{(1,14)} = 0.23$<br>$p = 0.64$<br>(ns) | $F_{(1,12)} = 0.52$<br>$p = 0.48$<br>(ns) | $F_{(1,14)} = 2.37$<br>$p = 0.15$<br>(ns) | - | $F_{(1,12)} = 3.03$<br>$p = 0.11$<br>(ns) |
| Basal blood glucose | $F_{(1,14)} = 16.6$<br>$p = 0.0011$<br>(**) | $F_{(1,14)} = 0.16$<br>$p = 0.69$<br>(ns) | $F_{(1,14)} = 33.6$<br>$p < 0.0001$<br>(****) | - | - | $F_{(1,14)} = 4.1 \times 10^{-4}$<br>$p = 0.98$<br>(ns) | $F_{(1,12)} = 0.015$<br>$p = 0.90$<br>(ns) | $F_{(1,14)} = 0.82$<br>$p = 0.38$<br>(ns) | - | $F_{(1,12)} = 0.15$<br>$p = 0.71$<br>(ns) |
| Peak size | $F_{(1,54)} = 0.20$<br>$p = 0.66$<br>(ns) | $F_{(1,54)} = 16.3$<br>$p = 0.0002$<br>(***) | $F_{(1,54)} = 7.26$<br>$p = 0.0094$<br>(**) | - | - | $F_{(1,54)} = 1.26$<br>$p = 0.27$<br>(ns) | $F_{(1,54)} = 1.26$<br>$p = 0.27$<br>(ns) | $F_{(1,54)} = 11.7$<br>$p = 0.0012$<br>(**) | - | $F_{(1,54)} = 2.30$<br>$p = 0.14$<br>(ns) |

**Supplementary table 3b:** Vanderbilt MMPC-Live C57BL/6J mice. C57BL/6J mice were given two oGTTs (MDA-oGTT and gavage-oGTT one week apart) in a randomised crossover. Glucose tolerance test (GTT) curve data for each sex were analysed by 3-way repeated measures analysis of variance (RM-ANOVA) for time, sex, and route of administration (RoA) interaction with Tukey’s multiple comparisons test. Baseline-corrected area under curve (AUC) and peak size data were analysed by 2-way RM-ANOVA for sex and RoA interaction. Bolded values indicate statistical significance (*p*<0.05).

| Vanderbilt C57BL/6J mice (Figure 3A-C; N = 16 [8 male and 8 female]) |  |  |  |  |  |  |
| --- | --- | --- | --- | --- | --- | --- |
|  | Sex | RoA | Time x RoA | Time x Sex | RoA x Sex | Time x RoA x Sex |
| C57BL/6J GTT curve | $F_{(1,14)} = 0.34$<br>$p = 0.57$<br>(ns) | $F_{(0.4,5.2)} = 20.4$<br>$p = 0.012$<br>(*) | $F_{(2.6,36.2)} = 7.76$<br>$p = 0.0007$<br>(***) | $F_{(6,84)} = 1.22$<br>$p = 0.30$<br>(ns) | $F_{(1,14)} = 2.86$<br>$p = 0.11$ (ns) | $F_{(6,84)} = 3.37$<br>$p = 0.005$<br>(**) |
| Baseline subtracted AUC | $F_{(1,14)} = 1.25$<br>$p = 0.28$<br>(ns) | $F_{(1,14)} = 18.9$<br>$p = 0.0007$<br>(***) | - | - | $F_{(1,14)} = 4.83$<br>$p = 0.045$<br>(*) | - |
| Peak size | $F_{(1,14)} = 8.12$<br>$p = 0.013$<br>(*) | $F_{(1,14)} = 9.21$<br>$p = 0.009$<br>(**) | - | - | $F_{(1,14)} = 2.67$<br>$p = 0.12$ (ns) | - |

